# The saccade-induced temporal attentional gradient: Duration compression of the post-saccadic second event

**DOI:** 10.1101/2023.10.28.564527

**Authors:** Lingyue Chen, Lukasz Grzeczkowski, Hermann J. Müller, Zhuanghua Shi

## Abstract

Rapid eye movements can distort our perception of time, as evidenced by the phenomenon of Chronostasis, where the first event following a saccade appears to last longer than it actually does. Despite extensive research on this phenomenon, the effect of saccades on sequential post-saccadic events, beyond the first one, has never been investigated. To address this, in the present study we compared the subjective estimates of the time of first and second post-saccadic events (with fixation conditions as controls). We found saccadic eye movements not only to affect the perceived duration of the first post-saccadic event (Chronostasis), but also that of the second event. Specifically, when the second event immediately following the first event, it was subjectively compressed. When the second event was used as the (constant) reference interval, Chronostasis was enhanced. Remarkably, the compression effect persisted even when potential “attentional-blink”-induced processes, that might affect timing at the transition from the first to the second event, were eliminated. To explain our findings, we propose that saccades induce a transient temporal attentional gradient that results in an overestimation of the first and an underestimation of the second interval when the two intervals are temporally (near-) contiguous.

## Introduction

Subjective time is sensitive to various types of contextual modulation. When we become engrossed in reading, time seems to fly by. Actions, such as pressing a key or catching a ball, can also affect the timing of subsequent events. One classic example of this is the stopped-clock illusion, also known as the Chronostasis illusion (Yarrow et al., 2001): when making an eye movement to look at a ticking clock, the second hand appears to stop momentarily before continuing to move. And the first second after the eye landing is perceived as longer than the following seconds (Knöll et al., 2013; Yarrow, 2010; Yarrow et al., 2001, 2004). In a typical Chronostasis study, participants are presented with a digital counter (initially set to ‘0’) in the visual periphery, to which they have to make a voluntary saccade. Immediately after the saccade lands on the counter, it begins counting upwards (‘1’, ‘2’, …), the duration of each digit that stays on representing an interval. The first duration, the test or ‘target’ interval, is variable, while the subsequent ‘reference’ interval is fixed. Participants judge whether the test interval is longer or shorter than the reference interval. One common account of the Chronostasis illusion is that saccades, or rapid eye movements, create retinal blur and active suppression degrade visual input, resulting in uncertainty as to the onset of an event (i.e., interval). The brain simply assumes that the post-saccadic image has remained constant throughout the saccade. This assumption causes our brain antedate the event onset to the saccade onset, giving rise to an apparent expansion of the event (Yarrow et al., 2004).

Actions can influence the perception of time not only by causing a time expansion, but also by causing compression. For instance, Morrone and colleagues (2005) observed that the duration of a short interval during a saccade is perceived as shorter than its actual duration. They attributed this to a slowdown of the neural clock while the saccade is executed, which is different from the explanation advanced by Yarrow and colleagues (2004). However, there is at present no consensus regarding a unified account of time expansion and time compression, despite several attempts to formulate such an account (e.g., Georg & Lappe, 2007; Knöll et al., 2013). Morrone and colleagues conceded that “Chronostasis may be related in some way to the compression and inversion effects reported here, but the connection [to saccade-induced compression] is not obvious” (Morrone et al., 2005, p. 953). To disentangle these apparently opposing effects, Knöll and colleagues (2013) systematically investigated the spatial-temporal topography of Chronostasis and found that, rather than being limited to target location of the saccade, overestimation also occurred for peri-saccadic events (with an onset from 100 ms before to 50 ms after the saccade) at the initial fixation location or positions midway on the saccadic path. Chronostasis could even be induced by a reduction of stimulus visibility (by means of a rapidly flippable mirror in front of participants’ right eye, mimicking the visual effect of saccadic movement) in the absence of a saccade. This led Knöll et al. to argue that overestimation of the duration during the perisaccadic period is “a passive result of how the time of a stimulus onset is predicted by the visual system in general” (Knöll et al., 2013, p. 64).

Most studies of Chronostasis have focused on low-level sensory mechanisms to account for (perisaccadic) Chronostasis, deliberately avoiding explanations in terms of attentional mechanisms (Yarrow, 2010; Yarrow et al., 2001). In part, this was based on finding that Chronostasis was little affected by whether or not participants had to make a voluntary shift of attention, in response to an arrow cue, to the target location before they executed the saccade (Yarrow, 2010; Yarrow et al., 2001). However, attention and motor action are tightly, and likely obligatorily, coupled (Deubel & Schneider, 1996; Nobre et al., 2010; Shepherd et al., 1986): attention is shifted toward the target (location) of the motor action *prior* to the actual commencement of the action, and this is the case with both saccadic eye movements (Deubel & Schneider, 1996; Posner, 1980; Shepherd et al., 1986) and manual pointing movements (Baldauf et al., 2006). Also, allocation of attention can lead to various time distortions. For instance, an attended event is perceived as lasting longer than an unattended event of the same duration (Enns et al., 1999; Tse et al., 2004). Also, a focally attended event is processed faster than unattended events in its vicinity, consistent with a spatial gradient of visual attention (C. J. Downing, 1988; Mangun & Hillyard, 1988). Furthermore, the first event or an oddball event that captures attention is perceived as longer than subsequent events (Kanai & Watanabe, 2006; Pariyadath & Eagleman, 2007; Rose & Summers, 1995). To examine the relationship between attention and saccade-induced Chronostasis, Georg and Lappe (2007) compared the location of stimulus presentation at the saccade landing position with that at the midway point on the saccadic trajectory. Choronstasis was found to be prominent only at the saccade landing position, where attention was assumed to be focused (cf. Deubel & Schneider, 1996), rather than at the midway position on the saccadic trajectory. At variance with this, Yarrow (2010) reported Chronostasis to be unchanged when observers judged the duration of letter probes that did not appear at the saccadic landing position. Also, systematically varying event onset (in addition to location: saccade start-, mid-, and end-position), Knöll et al. (2013) found Chronostasis to be evident for events that occurred as late as 50 ms after the offset of the saccade (with a similar profile for all positions).

While most studies on Chronostasis have focused on the time distortion of the first perisaccadic event, it remains unclear whether saccades can cause further distortion beyond this event. Analogously to the *spatial* gradient of attention (Mangun & Hillyard, 1988), goal-directed actions may induce a *temporal* gradient: allocating attentional resources to the first event may limit the amount of resources available for processing subsequent events, leading to an uneven distribution of attention over time. In some sense, this is similar to the classic ‘attentional-blink’ phenomenon: participants are typically poor at processing a (second) target that follows a first one with little delay (Duncan et al., 1997; Shapiro et al., 1997) – which is attributed to attentional, and/or working-memory, resources being engaged by the first target and so being unavailable for encoding the second target. Of note, though, the attentional blink reflects a limitation in target-related attentional selection, which in itself does not necessarily require an overt (motor) action (e.g., responding to target 1 may be verbal and delayed until after the end of the visual stream of target and interspersed non-target events). In Chronostasis experiments, blink-type processes may actually contribute to the overestimation of interval 1, by impacting the timing of interval 2, especially if the two intervals are temporally contiguous. Upon the signal indicating the end of interval 1 (and the start of interval 2), the timing of interval must be stopped and the recorded time (or some trace of the whole interval) stored in some sort of working memory for comparison with the second interval. This attention-demanding process may interfere with the timing of interval 2, for instance causing a loss of internal-clock ticks and thus leading to an underestimation of the second interval (thus contributing to the relative overestimation of interval 1). Another, on the surface similar but fundamentally different, conceptualization would be that the saccadic action (i.e., the eye movement to the to-be-timed event) creates a temporally uneven gradient of attention, where attention influences the gate capacity that pulses generated by the internal clock can pass through (see, e.g., the attentional gate theory by Zakay & Block, 1996a). Such a saccade-induced temporal gradient would then lead to an expansion of interval 1 (more clock ticks) and a compression of interval 2 (less clock ticks), both giving rise to an underestimation of the second event’s duration. Consequently, with both the attention-blink and the saccade-induced temporal-gradient accounts: if the reference interval for duration judgments is placed at the second temporal position, the underestimation may contribute to perisaccadic Chronostasis, i.e., the overestimation of the duration of the first event.

On this background, we conducted two experiments to investigate saccade-induced duration distortions that occur beyond the post-saccadic first interval, with a focus on the post-saccadic second interval. To ensure comparability with standard Chronostasis studies, we employed essentially the same paradigm, but critically, in Experiment 1, we varied the positioning of the reference interval, which could either immediately follow the test interval (No-Gap condition) or be delayed by two seconds (Gap condition). We expected Chronostasis to be increased when the reference follows the test interval without gap, which would be consistent with both accounts sketched above. In Experiment 2, we directly compared the first or, respectively, second interval after the saccade to a fixed reference interval temporally far from the saccade. Given that the reference interval is separated in time from both the first and the second test interval, the timing of the reference interval should not be affected by an ‘attentional blink’. However, if saccades induce an uneven temporal attentional gradient, the second post-saccadic event should be underestimated relative to the reference event – consistent with account 2 above.

## Experiment 1

### Methods

#### Participants

Twenty-one health participants, all with normal or corrected-to-normal vision, were recruited (mean age: 26.0 years; 9 females and 12 males). The sample size was determined based on previous studies (Morrone et al., 2005; Yarrow et al., 2001), which had an average of 16 (range: 4 to 30) participants. To have sufficient power with a similar design, the sample size was increased to 21 participants. Participants were not aware of the purpose of the experiment. The study, including Experiment 1, was approved by the Ethics Committee of the LMU-Munich Faculty of Psychology and Pedagogics. Participants provided informed consent prior to the experiment, and were compensated at a rate of 9€ per hour for their service.

#### Apparatus

The experiment was conducted in a quiet and dark laboratory cabin. Participants sat in front of a display monitor (ViewPixx LCD, VPixx Technologies Inc.; screen refresh rate: 120 Hz), with a viewing distance of 60 cm maintained by a chinrest. Their eye movements were tracked and recorded by an EyeLink 1000 system (SR Research Ltd.), with a sampling rate of 1000 Hz. Behavioral responses were collected via a standard keyboard. The experimental program was coded in Matlab with the PsychToolbox (Brainard, 1997; Pelli, 1997) and the Eyelink toolbox (Cornelissen, Peters, & Palmer, 2002).

#### Stimuli and procedure

There were two saccade and two fixation conditions (see Fig 1). In the saccade conditions, participants had to make a voluntary saccade, from a central white fixation dot (size: 0.1° of visual angle; luminance: 68 cd/m^2^), toward one of two possible locations indicated by two white disks (1°, 68 cd/m^2^). The disks were located diagonally opposite relative to the display center (randomly, either one left-down and the other right-up or, respectively, one left-up and the other right-down, at eccentricity of 10°), and which of the two disks the (voluntary) saccade was made to on a given trial was chosen by the participant. A trial started with the central fixation marker (on a dark gray background, 5 cd/m^2^), prompting the participant to fixate the marker for 1 s (with a spatial-error tolerance of ±2° of visual angle). The dot then changed into green, providing the cue to execute the saccade. Once the eyes landed on the target dot, the irrelevant disk on the opposite side was extinguished immediately to minimize any potential distractions. At the chosen location, a green concentric ring around the target disk (30 cd/m^2^) was flashed for 25 ms, indicating the start of the test interval. Following a randomly varying interval (duration selected from 125, 250, 375, 500, 625, 750, and 875 ms), a second flash (of a green ring around the disk) marked the end of the test interval. In the Saccade/No-gap condition (Fig 1, top row), the flash marking the offset of the test interval also indicated the onset of the (fixed-length) reference interval. In the Saccade/Gap condition, by contrast, a third flash was presented after a constant, 2000-ms gap to mark the onset of the reference interval (Fig 1, third row). In all conditions, the final flash indicated the end of the reference interval. Then, after a blank interval of 500 ms, participants were prompted with the question displayed with “which interval lasted longer: the first or the second?”. Participants had to make a two-alternative forced-choice (2AFC) by pressing the left or right arrow key for the first or, respectively, the second interval as having been perceived as longer. Each test interval was repeated 20 times, in random order with the other test intervals.

**Fig 1.**
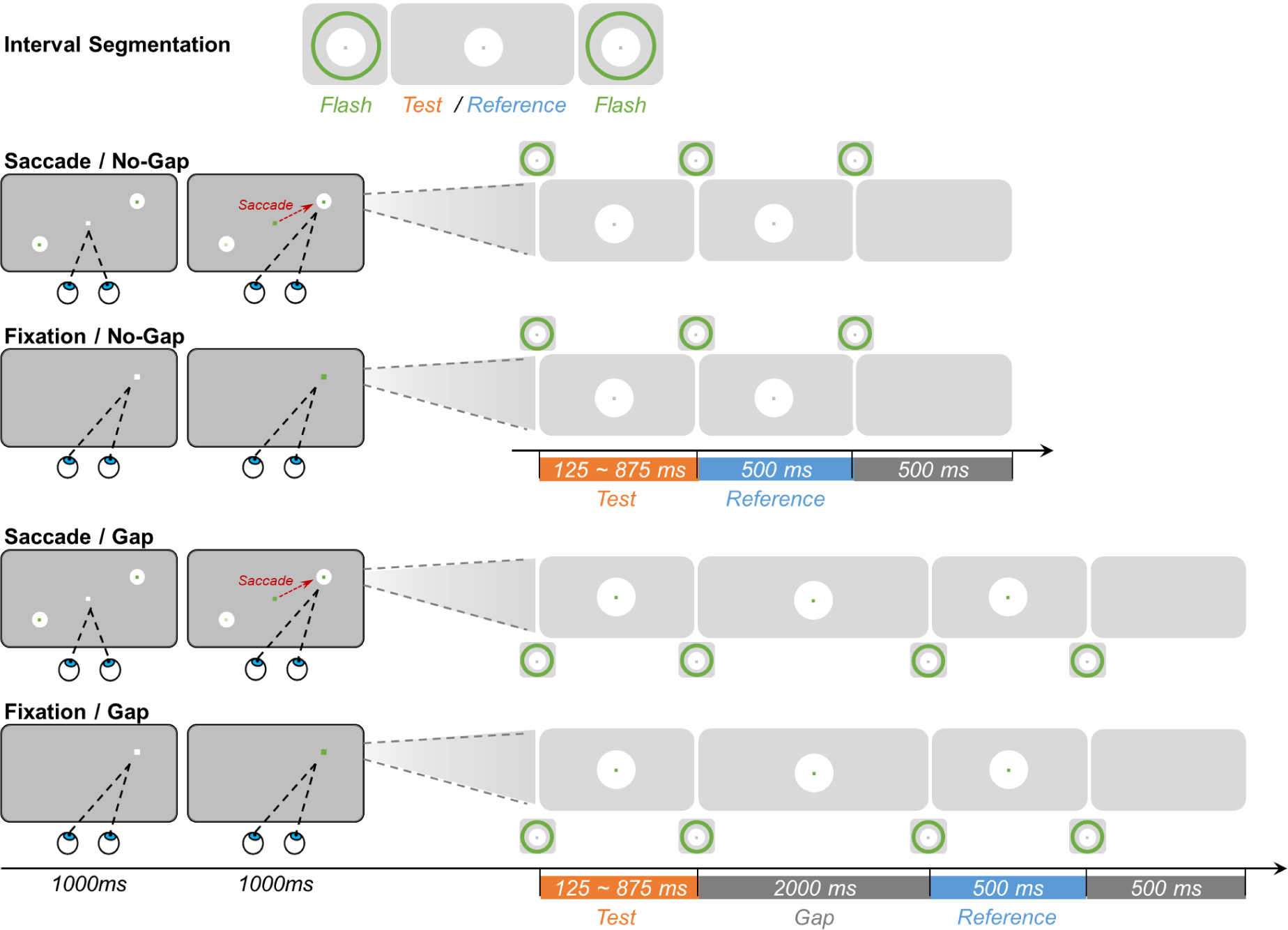
Experiment 1 comprised four blocked conditions: Saccade/No-gap, Fixation/No-gap, Saccade/Gap, and Fixation/Gap. The top row shows the interval segmentation for all conditions. In the bottom four rows, the left panels show the displays for the eye-movement phase, where each trial started (separately for the Saccade and Fixation conditions); and the right panel shows the subsequent stimulus changes in central vision for the interval-comparison phase (separately for the Gap and No-Gap conditions). In the Saccade conditions (1st and 3rd row), participants fixated a central white marker dot, which (after 1000 ms) turned green, cueing a saccade to one of the two peripheral disk locations. Landing on the saccade target triggered the interval-comparison phase. In the Fixation conditions (2nd and 4th row), participants fixated the dot in one corner of the screen, which turned green after one second. In the interval-comparison phase (right panels), brief (25 ms) flashes of concentric green rings demarcated the test and reference intervals. In the No-Gap conditions (upper two rows), three consecutive flashes demarcated the test and reference intervals, whereas in the Gap conditions (lower two rows), four flashes demarcated the test, a gap, and the reference interval, respectively. Participants indicated which interval (the test or the reference interval) was longer by pressing a corresponding button.

To distinguish the time distortion induced by saccadic action, two baseline Fixation control conditions were introduced, one with and the other without a gap (as shown in the second and fourth row of Fig 1). In these conditions, participants were required to fixate a single dot without making any eye movements. The position of the dot was randomly chosen from the (four) possible landing locations in the saccade conditions. Participants were instructed to maintain fixation on this location throughout the entire trial. After one second, the color of the dot changed from white to green, indicating that the test and reference intervals would ‘soon’ be presented. Similar to the Saccade conditions, a white disk appeared (with the green dot staying on in the center) after 1000 ms, roughly matching the time taken for selecting the target disk and making a saccade to it in the Saccade conditions. The subsequent sequence of events was then the same as in the saccadic conditions.

During each trial, participants’ eye movements were monitored. In the Fixation conditions, they had to keep their gaze within a specific area around the fixation marker (spatial-error tolerance of ±2° of radius) for the entire duration of the trial. In the Saccade conditions, they had to make the correct saccade towards the target and then maintain fixation within the designated area. Trials on which the participant blinked or fixated outside the designated area were considered invalid and immediately terminated, accompanied by a warning beep (5000 Hz, 31 Db) for 100 ms. Such failed trials (which occurred, on average, in 9.18%) were randomly retested at the end of each block to ensure that all conditions had an equal number of 140 valid trials.

Together, the experiment included four combinations of conditions, based on two actions (Saccade vs. Fixation) and two reference types (No-gap vs. Gap). These conditions were tested in blocks, with the order of four blocks randomly assigned to each participant but counterbalanced across participants. Henceforth, the four conditions will be referred to as Saccade/Gap, Saccade/No-gap, Fixation/Gap, and Fixation/No-gap, respectively. The entire experiment lasted approximately two hours, with participants taking breaks between blocks as needed.

Prior to the formal experiment, participants completed two training blocks (Saccade/Gap and Fixation/No-gap conditions) to become familiar with the tasks. Each training block included 20 test intervals (100 and 1000 ms, not included in the formal test), with the standard reference interval of 500 ms. To help participants understand the task, accuracy feedback was provided at the end of each trial, in the form of a warning beep (2000 Hz, 43 Db, 100 ms) upon an incorrect response (no such accuracy feedback was provided during the format test session). The formal experiment started when the accuracy rate was above 80%, otherwise, an additional round of training was added.^1^

#### Data analysis

The ‘First’ vs. ‘Second’ responses (to the question which of the two intervals was longer) were transformed into ‘Longer’ vs. ‘Shorter’ judgments of the test interval relative to the reference interval. The mean proportion of ‘Long’ responses for each test interval was then calculated for each condition. Psychometric curves were estimated using the R package QuickPsy (Linares & López-Moliner, 2016) for each participant in each condition, with lapse and guess rates taken into account (the mean estimated lapse rate was 0.1 in Experiment 1, which was then taken as a reference for Experiment 2). The psychometric curves allowed us to obtain two key parameters: the point of subjective equality (PSE) and the just-noticeable difference (JND). The PSE indicates the transition threshold between short and long judgments, while the JND provides an index of temporal discrimination sensitivity. Finally, these parameters were examined in repeated-measures analyses of variance (ANOVAs), conducted with the factors Reference Timing (with vs. without Gap) and Action (Saccade vs. Fixation).

## Results and Discussion

Experiment 1 examined whether presentation of the fixed reference interval immediately following (vs. following with a delay) the variable test interval would enhance Chronostasis. Fig 2A depicts the psychometric curves for one typical participant, with each curve representing the ratio of “long test interval” responses relative to the reference. A lower point of subjective equality (PSE) than the actual reference duration (500 ms) indicates an overestimation of the test interval, meaning that a shorter interval would be required to match the standard. The mean PSEs (with associated standard errors, ± SE) were 415 (± 21), 460 (± 14), 441 (± 18), and 496 (± 15) ms for the Saccade/No-gap, Saccade/Gap, Fixation/No-gap, and Fixation/Gap condition, respectively (Fig 2B).

**Fig 2.**
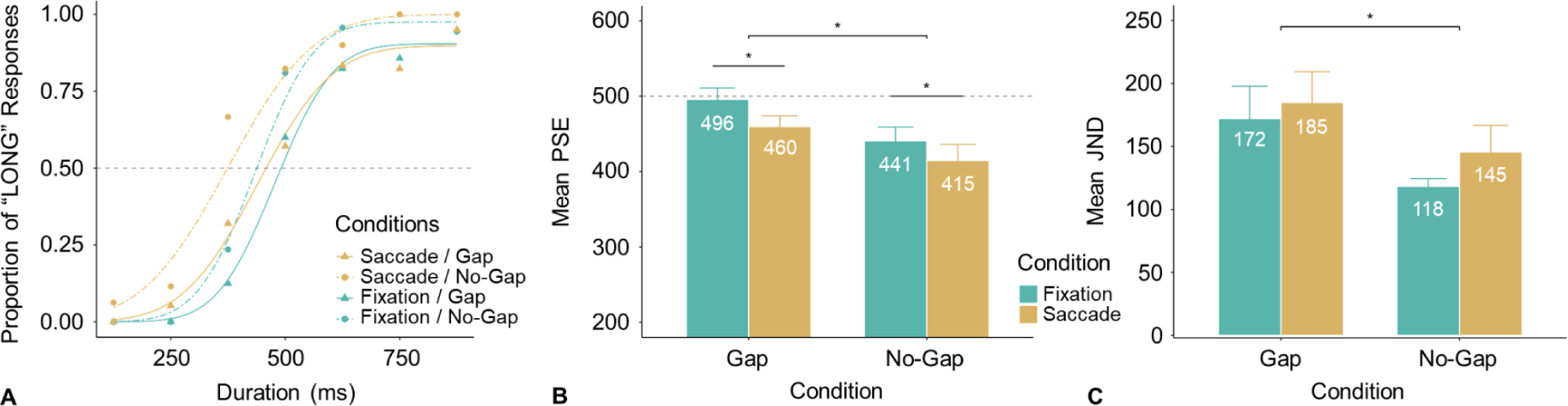
Results from Experiment 1. (**A**) A typical example of behavioral responses (dots) and fitted psychometric curves from one participant. The mean PSEs (**B**) and JNDs (**C**) are shown for the four conditions, across all participants. The dashed horizontal line marks the reference interval (500 ms). The smaller the PSE, the more the dilation on the test interval, as it would require a shorter (test) duration to be perceived as long as the standard (reference) duration. (*: p < .05).

The difference between the Saccade and Fixation conditions was significant, *F*(1, 20) = 5.64, *p* = .028, 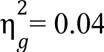, with a reduction of 31 ms for the Saccade conditions, evidencing the Chronostasis (Yarrow et al., 2001). The main effect of Reference Timing was significant, *F*(1, 20) = 6.41, *p* = .020, 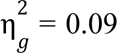: the test interval was perceived as longer when the reference immediately followed the test interval (No-gap condition), compared to when there was a 2-second gap of between the two intervals (Gap condition). The Action × Reference Timing interaction was non-significant, *F* (1, 20) = 0.15, p = .703, 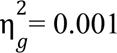.

For follow-up analysis, we conducted *t*-tests (two-tailed, adjusted for multiple comparisons) to examine whether the PSEs were smaller than the actual, 500-ms reference. The results revealed the PSEs to be significantly smaller in the Saccade/Gap condition (*t*(20) = −2.83, *p* = .014) and in both No-Gap conditions (Fixation: *t*(20) = −3.25, *p* = .008, Saccade: *t*(20) = −4.04, *p* = .003); however, the PSE was close to the actual 500 ms in the Fixation/Gap condition (*t*(20) = −0.28, *p* = .784). These results are indicative of a general tendency to (relatively) underestimate the second (i.e., reference) interval when it immediately follows the first (i.e., test) test interval.

The mean JNDs (± SE) were 145 ± 21, 185 ± 24, 118 ± 6, and 172 ± 26 ms for for the Saccade/No-gap, Saccade/Gap, Fixation/No-gap, and Fixation/Gap condition, respectively (Fig 2C). A two-way repeated-measures ANOVA revealed the main effect of Reference Timing to be significant, *F* (1, 20) = 4.59, *p* = .045, 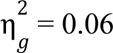; the main effect of Action, *F* (1, 20) = 2.47, p = .132, 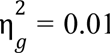, and the Action × Reference Timing interaction, *F* (1, 20) = 0.20, *p* = .660, 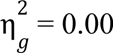, were non-significant. Introducing a gap induced additional memory decay, worsening temporal discrimination sensitivity.

In short, Experiment 1 revealed a significant Chronostasis effect (31ms), alongside a significant impact of the temporal positioning of the reference interval (50 ms). Specifically, when the reference interval immediately followed the test interval, the test interval was perceived as longer than when the reference was presented after a 2-second gap. Looked at in terms of the second (reference) event, the reference interval was ‘compressed’ relative to the first interval. While this ‘compression’ effect appears in part linked to the action, as evidenced by the lower PSE with vs. without a saccade (415 ms vs. 441 ms), the main source of the ‘compression’ with two contiguous intervals may be an ‘attentional blink’, potentially exacerbated by the intervals being defined by abrupt-onset flashes, i.e.: salient stimuli that engage attention automatically (Remington et al., 1992; Yantis & Jonides, 1990). That is, reinforced by the first flash, attention is allocated to and engaged by the first interval, causing a blink-type difficulty with starting the timing of the second interval, in particular as the second flash, demarcating the transition from the first to the second interval, prompts (executive) processes directed to the first interval. Attending to the second interval may therefore be delayed (by an ‘attentional blink’), causing the second interval to be underestimated (see also the ‘attentional-gate’ account of (Zakay & Block, 1996b). In contrast, when the reference appeared two seconds after the end of the first interval, attention can be reallocated to the reference event without difficulty. This avoids the attentional blink, leaving a minor saccade-induced Chronostasis effect (36 ms, based on the PSEs of 496 ms vs. 460 ms in the fixation vs. saccade conditions).

To investigate the potential contribution of an attentional-blink effect, we conducted a control experiment without any eye-movement ‘action’ (i.e., the stimuli were presented at central fixation; see Appendix A for details). The results showed that the abrupt flash onset indeed significantly induced an attentional blink, causing the second event to be perceived as shorter than the first event (PSE of 389, as compared to the fixed 500-ms test interval). This flash-induced attentional blink might potentially overlay any *action*-induced time distortions. When introducing conditions (in the control experiment) in which the test and reference intervals were defined by color identity, the two consecutive color-defined intervals were perceived as similar in duration: the PSE for the test interval was 494 ms, which is close to the actual (500-ms) duration of the reference interval. We attribute this to the change from one to the other color generating less visual onset transients (compared to the abrupt-onset flashes) and so less confounding by blink-type processes. For this reason, we used color identity to define the interval duration in Experiment 2.

## Experiment 2

Experiment 2 was designed to directly measure action-induced distortions to the second vs. first interval following the saccadic action, with the reference interval presented 1.2 seconds after the second interval. Assuming saccades induce an uneven temporal distribution of attention like the attentional blink, we expected the second post-saccadic event would be underestimated relative to the reference event.

### Method

#### Participants

21 new participants (mean age: 26.7 years; 11 females and 10 males) were recruited for Experiment 2. All had normal or corrected to normal vision and color vision, and were naïve as to the purpose of the experiment. Before starting the experiment, participants provided informed consent. Payment was again at a rate of 9 € per hour.

#### Stimuli and procedure

The setup as in Experiment 2 was similar to that in Experiment 1, with the following differences. The saccadic (target) disks were positioned 6° to the left and right of the central fixation point. Each trial started with the central fixation marker presented for 500 ms, followed by a white arrow cue (< or >) indicating the location of the target on a given trial. The target and non-target disks both (i.e., as a group) changed from white to green (or, respectively, from white to red) immediately after the saccade offset, marking the onset of the first interval. At a designated time, the disk changed to red (or, respectively, green), signaling the end of the first interval and the start of the second interval. Following another designated time (see details below), the disk turned white for 1200 ms to create a gap before the reference interval. The disk then changed color to either green or red marking the onset of the reference, depending on the selected test interval, which was blocked per ‘session’ (i.e., either the first interval was consistently the test interval in a session, or the second interval). Participants had to compare the test interval, either the first or second interval (which shared the same color with the reference interval, red or green) with the reference interval and judge which one was longer, by pressing the left or right arrow key.

In Experiment 2, the PSEs (and JNDs) were measured for two post-saccadic intervals: the first interval and the second intervals (hereafter labeled S1 and S2, where S denotes the Saccadic conditions). The test and the reference intervals were assigned the same color, while the non-test interval used a different color. The colors were isoluminant and counterbalanced assigned to the first and second intervals across participants (i.e., half of the participants saw green and red as the first and second intervals in all conditions, while the other half saw red and green respectively). The test interval varied randomly from 150 to 1050 ms in increments of 150 ms (7 levels), while the non-test interval and the reference were set to 600 ms^2^ (see Fig 3).

**Fig 3.**
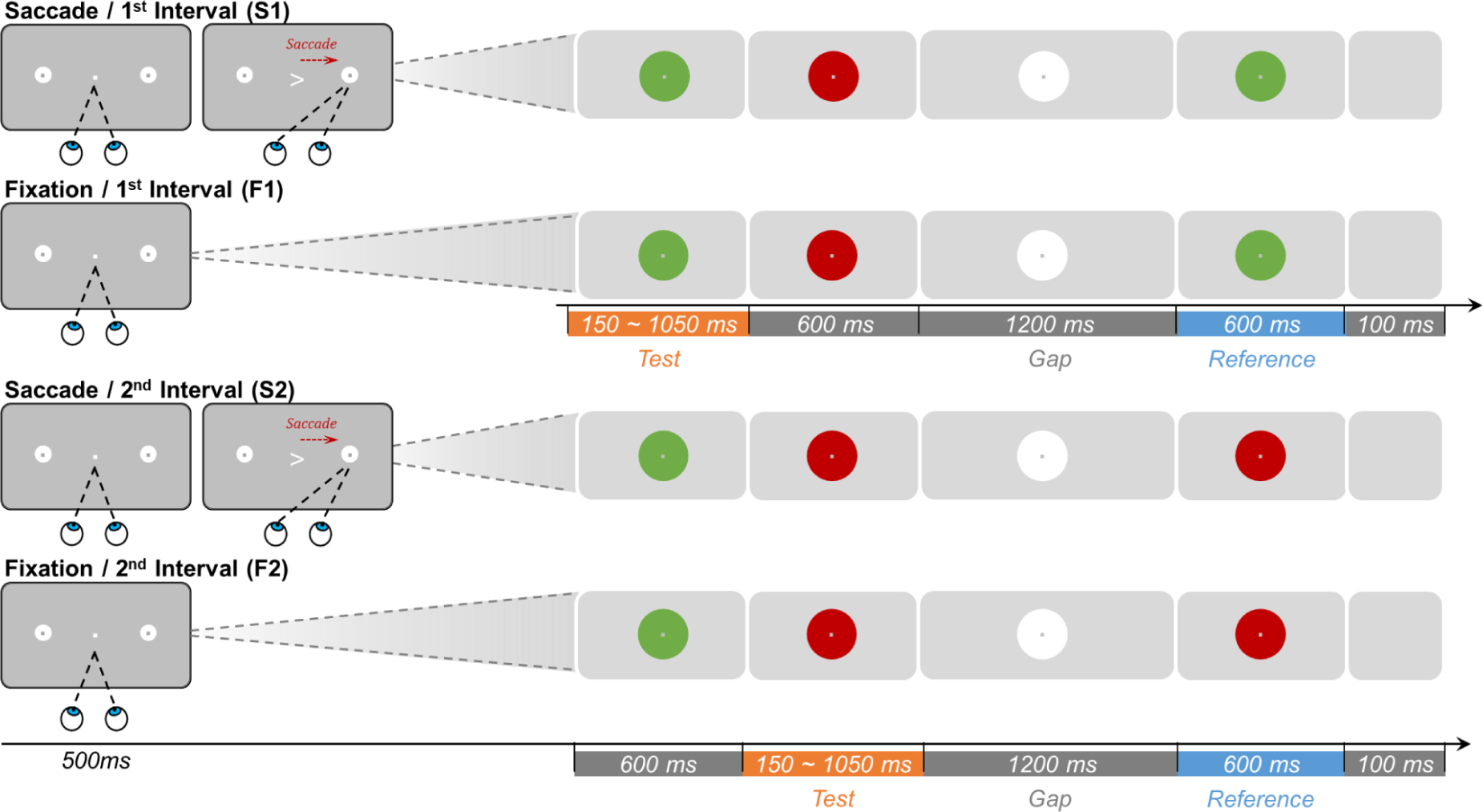
Schematic illustration of the procedure of Experiment 2. There were four blocked conditions: Saccade/1st Interval (S1), Fixation/1st Interval (F1), Saccade/2nd Interval (S2), and Fixation/2nd Interval (F2). The left panels illustrate the displays for the trial-initial action phase, separately for the Saccade and Fixation (action) conditions. The right panels depict the subsequent stimulus changes for the phase of interval comparison. In the Saccade conditions, participants fixated the central fixation dot for 500 ms, whereupon they received a spatial arrow cue prompting them to make a saccade to the indicated target disk (1st and 3rd rows); landing on the saccade target then triggered the interval-comparison phase. In the Fixation conditions, there was no central change, requiring participants to maintain fixation in the center (2nd and 4th rows), and the interval-comparison phase began automatically after 500 ms of fixation. In the interval-comparison phase, the intervals were demarcated by changes of the disk color. In the S1 and F1 conditions (upper two rows), the first interval shared the same color with the reference interval, and so interval 1 was the task-relevant test interval (blocked per S1 and F1 session). In the S2 and F2 conditions (bottom two rows), the second interval shared the same color with the reference interval, and so interval 2 was the task-relevant test interval (blocked per S2 and F2 session). The color of the reference interval varied across conditions (green or red), while the color of the gap interval was always white. Participants indicated which interval – the test or the reference interval – was longer by pressing a corresponding key.

There were also two analogous baseline conditions without eye movements, one in which the first interval was the test interval and one in which the second interval was the test interval (hereafter labeled F1 and F2, where F denotes the Fixation conditions; see Fig. 3, second and fourth rows). The procedure was identical to the saccade sessions (S1 and S2), except that observers were asked to maintain fixation on the central dot throughout the trial. After 500 ms fixation, the first two intervals and the reference were presented in the same manner as in the saccade sessions.

Each participant completed all four experimental conditions (S1, S2, F1, and F2) in a random order, with condition order counterbalanced across participants. Each session consisted of seven intervals that were randomly repeated 20 times, 10 per each side. As in Experiment 1, participants’ eye-movements were monitored throughout each trial. Any trials with incorrect eye movements (on average, 12.77%) were retested in a random order at the end of each block, ensuring that all conditions had an equal number of 140 valid trials.

## Results and Discussion

Fig. 4A depicts typical responses from one participant and associated fitted response curves, while Fig.s 4B and C show the average PSEs and JNDs across all participants.

**Fig 4.**
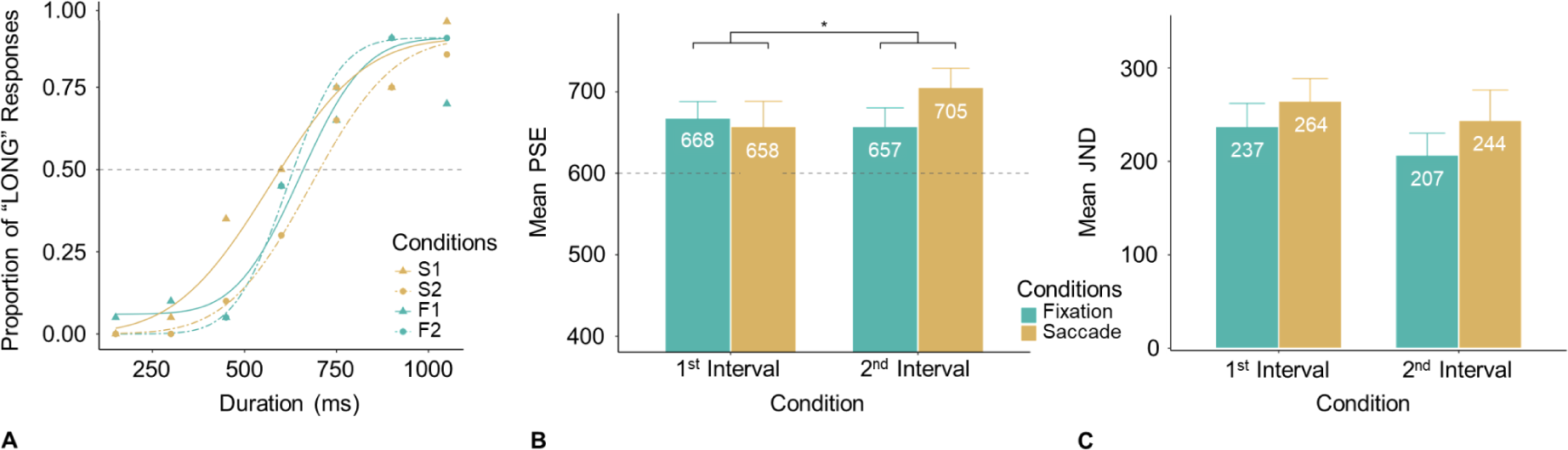
Results from Experiment 2. A typical example of behavioral responses (dots) and fitted psychometric curves from one participant (**A**). The mean PSEs (**B**) and JNDs (**C**) for the four conditions from all participants. The dashed horizontal line marks the reference interval (600 ms). The increased greater PSE value for the second test interval in the saccade condition (compared to the other conditions) indicates a compression of this interval, as it would require a longer duration for it to be perceived as long as the delayed reference interval. (*: p < .05).

The mean PSEs (± SE) for all participants were 658 ± 31, 705 ± 23, 668 ± 20, and 657 ± 23 ms for the S1, S2, F1, and F2 conditions, respectively. A repeated-measures ANOVA revealed only the Action (Saccade, Fixation) × Test-Interval (1, 2) interaction to be significant, *F*(1, 20) = 5.84, *p* = .025, 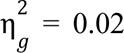; the main effects were non-significant: Action, *F*(1, 20) = 1.09, *p* = .309, 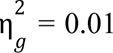; Test-Interval, *F*(1, 20) = 1.03, *p* = .323, 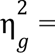 0.01. The interaction was largely due to the ‘odd-one-out’ large PSE in the S2 condition, relative to comparable PSEs in the other three conditions. Further post-hoc comparisons revealed a significant difference between S2 and S1 (*t*(20) = 2.00, *p* = .045, *Cohen’s d* = 0.45) and between the S2 and F2 condition (*t*(20) = 2.12, *p* = .045, *Cohen’s d* = 0.47), but not between F1 and S1 (*t*(20) = −0.49, *p* = .627, *Cohen’s d* = −0.11). These findings confirm that the post-saccadic second interval (S2) was greatly compressed. However, we failed to show any significant Chronostasis. Given that the reference interval was constant (600 ms), we further conducted simple *t*-tests to examine the absolute over-or underestimates. The results showed that the PSEs were larger than 600 ms (*p*s < .05). That is, regardless of the presence of a saccade, the *first* interval (F1 or S1) was perceived as shorter than the reference interval, even though the reference interval was separated from the end of the first interval by 1800 ms (which is similar to the 2000-ms separation in Experiment 1). In other words, when the first interval was judgment-relevant, it too was compressed (relative to the delayed reference interval) to some extent. We tentatively attribute this surprising finding to the presence of a second (irrelevant) interval intervening between the task-relevant first and the reference interval, which may have given rise to a recency effect for the last, reference interval (for more detailed arguments, see the General Discussion).

The mean JNDs (± SE) were 264 ± 24, 244 ± 33, 237 ± 25, and 207 ± 23 ms for the S1, S2, F1, and F2 conditions, respectively (Fig 7C). A two-way repeated-measures ANOVA revealed no significant effects (Action, *F*(1, 20) = 3.11, *p* = .093, 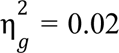; Test-Interval, *F*(1, 20) = 1.91, p = .183, 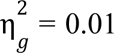; interaction, *F*(1, 20) = 0.05, p = .818, 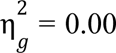, was significant.

## General Discussion

The subjective perception of time can be distorted by actions (Merchant & Yarrow, 2016). Previous studies have shown that saccadic actions can either expand (Morrone et al., 2005; Yarrow et al., 2001)) or compress (Morrone et al., 2005; Yarrow et al., 2001) the perceived duration of the peri-saccadic first event. In the present study, we investigated how making a goal-directed saccade influences the duration perception of subsequent events, particularly of the post-saccadic second interval, which is typically used as a reference interval in studies of Chronostasis (e.g., Georg & Lappe, 2007; Park et al., 2003; Yarrow et al., 2001, 2004).

### Summary of Experiments and Critical Findings

Experiment 1 showed a greater Chronostasis effect when the reference interval immediately followed the post-saccadic first (i.e., test) interval (i.e., No-Gap) as compared to when there was a 2-second gap between the two intervals. We considered two possible explanations for this overestimation of the post-saccadic time under No-Gap conditions. First, the post-saccadic interval may have been expanded by attention being shifted to this event, owing to the inherent coupling of spatial attention and saccadic eye movements (Deubel & Schneider, 1996; Shepherd et al., 1986). This would have resulted in a transient post-saccadic attentional enhancement of the first event and consequently a decline of attention to the (immediately following) second event, i.e., a ‘compression’ of the post-saccadic second interval relative to the first interval. Second, attention may have been engaged by the first event, especially because this event was signaled by an abrupt-onset flash ‘capturing’ attention (Remington et al., 1992; Yantis & Jonides, 1990). This may have given rise to an ‘attentional blink’ at the end of the first interval, a control-demanding point (Kawahara et al., 2006) at which the ‘clock’ had to be stopped and the result buffered in working memory. These processes could have delayed the timing of – and thus effectively ‘compressed’ – the immediately following reference interval. In contrast, when the reference followed the target interval after a 2-second gap, the blink would have occurred during the gap and so have left the timing of the second interval unaffected. Of note, an ‘attentional-blink’-type compression of the second (reference) interval could only explain part of the relative overestimation of the first (test) interval, as there remained a Chronostasis effect in the No-Gap saccade (vs. fixation) condition. A control experiment without eye movements (reported in the Appendix) confirmed that a blink-type effect was evident when the intervals were demarcated by salient abrupt-onset flashes, but not when they were demarcated by (isoluminant) color changes.

Given this, Experiment 2 was designed to more directly examine the hypothesis that saccadic actions induce a temporal attentional gradient that transiently enhances the timing of the first post-saccadic event, while (as the initial boost fades out) degrading the timing of the immediately following, second event. To investigate this, we introduced several changes to the experimental set-up compared to Experiment 1: Both the first and second intervals were test intervals – though blocked per condition, so participants knew which interval was task-relevant and to be attended and which one could be ignored. Additionally, the reference interval was presented with a gap after the end of the second interval, and the intervals were marked by color changes rather than abrupt-onset flashes. The latter measures were implemented to minimize blink-induced distortions and so to isolate any temporal-gradient effects. According to the “temporal attentional-gradient” account, the second test interval should be compressed (relative to the reference interval), while the first interval should be expanded.

The results showed that the perceived duration of the first interval was similar in the Saccade to the Fixation condition, failing to show significant saccade-induced Chronostasis. In fact, if anything, the first interval was underestimated (“compressed”) compared to the reference duration, in both conditions. This is a finding that does not easily square with the “temporal attentional-gradient” account, requiring further discussion (see below).^3^ Crucially, however, when the second interval was the test interval, the saccade caused a significant compression as compared to the fixation control condition, as predicted by the “temporal attentional-gradient” account.

One possibility why we found no saccade-induced Chronostasis effect for the first interval may have to do with the presence of a second (irrelevant) interval intervening between the task-relevant first and the reference interval. This may give rise to a recency effect for the last – the reference – interval, manifesting in a degree of ‘compression’ of the preceding intervals (which suffer from ‘trace decay’). Consistent with this would be the finding that the underestimation of the first interval was similar to that of the second interval in the fixation condition. Alternatively, it is known that a group of intervals can be assimilated to the ensemble mean at low-level perceptual processing (Baykan et al., 2023; Burr et al., 2013; Nakajima et al., 1992; Ren et al., 2020). So, the significant compression of the second interval might assimilate the first interval, negating any minor Chronostasis that occurred to the first interval (as shown in Experiment 1). Also, as shown by Knöll et al. (2013), the Chronostasis effect can quickly disappear when the critical event is presented 50 ms after the saccade. These factors may all have contributed to the lack of saccade-induced Chronostasis in Experiment 2.

### Theoretical considerations

The present study shed light on the intricate nature of subjective time distortions induced by saccades. Our results indicate that the stimulus onset, saccadic action, and the timing of the reference interval all play crucial roles in duration judgments. In Chronostasis experiments, a digital clock is often used to display a sequence of time intervals demarcated by the clock changing digits. Typically, the test interval is the first digit flip (from 0 to 1) that occurs after the saccade, while the reference interval is the interval immediately following the test interval – which is comparable to the No-Gap condition in our Experiment 1 (Georg & Lappe, 2007; Park et al., 2003; Yarrow et al., 2001). However, as we demonstrated in Experiment 2, the immediately following reference itself (i.e., the second interval) can be impacted by the saccade, as saccade-coupled attention may create a temporal attentional gradient that boosts processing temporarily in the peri-saccadic period, but compromises the processing of immediately following events. The ensuing compression of the post-saccadic second interval was evident in Experiment 2 when it was directly compared to the reference interval presented after a 1.2-seconds gap. Recall that Experiment 2 had two fixation baselines, which both produced comparable PSEs (668 ms and 657 ms for the first and the second interval, respectively). This suggests that without any eye movements, the two intervals were perceived similarly (albeit shorter than the reference interval, which followed the second interval after a gap). Therefore, the greatly increased compression of the post-saccadic second interval (PSE of 705 ms) could only have been caused by the preceding saccadic eye movement. Unlike the post-saccadic first interval, whose onset was highly uncertain due to the saccade, the onset of the second interval occurred well beyond the peri-saccadic time window (600 ms after the saccade). Accordingly, active compensation for the stimulus onset (Yarrow et al., 2001) or low-level sensory factors (Knöll et al., 2013), which have been proposed to account for Chronostasis, cannot readily explain the compression of the post-saccadic second interval.

As a limitation, we note that the compression of the post-saccadic second interval only contributes to (rather than fully explains) the Chronostasis effect, as previous studies have reported robust Chronostasis illusions when the reference was temporally further removed from the saccadic action, separated by a gap of 500 ms or even 1000 ms (e.g., Knöll et al., 2013; Yarrow et al., 2004).

One possible explanation for the compression of the post-saccadic second interval is an uneven spatio-temporal attentional gradient tied to the saccadic eye movement. Spatially, attention is concentrated on the landing position of the saccade (i.e., the saccadic target) and decreases from there gradually (Mangun & Hillyard, 1988). This gradient also accounts for the line-motion illusion (Downing & Treisman, 1997; Hikosaka et al., 1993), in which a flash preceding the onset of a closeby line leads to subjective motion of the line outwards from the position of the flash. Temporally, planning and executing a voluntary eye movement to a target location is coupled with an attention shift to the saccade target (e.g., Deubel & Schneider, 1996; Shepherd et al., 1986), giving rise to a relatively transient post-saccadic attentional enhancement of objects or events at this location (see also Müller & Rabbitt, 1989). Thus, the first post-saccadic event occurring there would benefit from this enhancement, while the second event would fall into a trough (perhaps analogous to the “inhibition-of-return” effect in the spatial domain; e.g., Klein & Ivanoff, 2008, for a review), compromising its temporal processing and leading to it being perceived as shorter than its actual duration.

Overall, this account is consistent with previous findings that attention modulates the Chronostasis effect. For example, Chronostasis was diminished when the peri-saccadic event was presented spatially outside the focus of attention, at a midway position on the saccadic trajectory (Georg & Lappe, 2007; but see Knöll et al., 2013). Here, we find that a saccade-induced temporal attentional modulation extends beyond the post-saccadic first event. Future work is required to substantiate this account. For instance, future studies could vary the onsets of the first and second second intervals to track the duration of the transient boost and the attentional trough induced by saccades. Further, investigating phase oscillations in the electroencephalogram, which have been linked to attentional blink (Zauner et al., 2012) and temporal expectation (Cravo et al., 2013; Nobre & Van Ede, 2018), might shed light on neural mechanisms underlying the modulation of post-saccadic time estimation.

To sum up, saccadic eye movements affect not only the perceived duration of the first post-saccadic event (Chronostasis) but also the subsequent events. Our findings indicate that the second post-saccadic event following immediately upon the first event is subjectively compressed – enhancing the Chronostasis effect when the second event is used as the reference interval. This compression is demonstrable even when ‘attentional-blink’-type distortions are minimized or eliminated. We propose that saccades induce a transient temporal attentional gradient, resulting in an overestimation of the first interval and an underestimation of the second interval immediately after the saccade.

## Appendix A. Baseline Control Experiment

## Baseline Comparison

According to the findings of Experiment 1, the flashes could have created abrupt changes that captured attention involuntarily, which could have led to a temporary inability to perceive the second event (also known as attentional blink). As a result, the second event was perceived as shorter than the first event. Given that our brain appears to prioritize attending to new object onsets over feature changes of existing objects (Yantis & Jonides, 1990), using color feature changes may generate less attentional capture than abrupt flash onsets. Therefore, in this control experiment, we compared the perceived duration between intervals defined by color changes vs. intervals defined by ring flashes.

### Method

#### Participants

20 new healthy participants, with normal color vision, were recruited (mean age of 27.1 years; 12 females and 8 males). All participants were naïve as to the purpose of the experiment. They provided informed consent prior to testing, and were remunerated at a rate of 9 € per hour.

#### Apparatus

The experiment was conducted in the same experimental cabin using the same hardware without Eyelink.

#### Stimuli and procedure

**Fig. A1.**
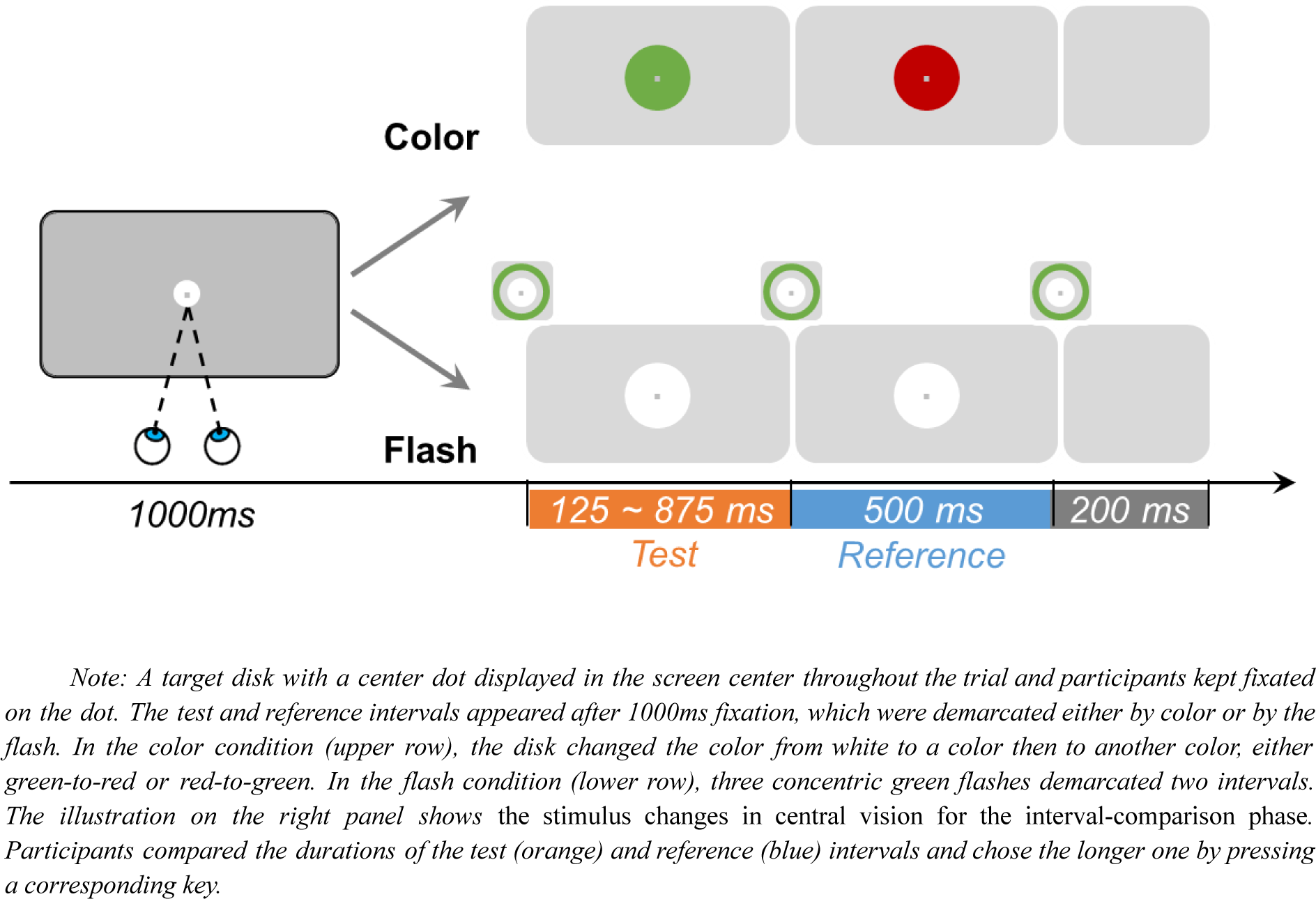
Schematic illustration of the procedure.

The conditions in this control experiment were essentially the same as in the No-Gap/Fixation condition of Experiment 1, with the following differences. Given that there was no saccade conditions, the white disk and fixation dot were presented in the screen center. Participants were instructed to fixate the center dot throughout the trial. There were two conditions, Flash vs. Color (Fig. A1, right bottom vs. upper panel), which were tested in separate blocks, with the order counterbalanced across participants. In the color condition, the start of the first – ‘test’ – interval was signaled by the central disk changing color (from white) to either green (for one half of the trials) or red (for the other half). The end of this interval and the start of the second – “reference’ – interval was signaled by another color change, from red to green or, respectively, green to red (where the colors assigned to the test and reference intervals was randomized across trials). The seven test intervals were randomly repeated 20 times each for both the color and flash blocks, resulting in 140 trials per block and 280 trials in total. Before formal testing in each (Flash, Color) condition, participants underwent a brief training block, in which the test durations were either 100 or 900 ms (presented in random order). Participants received accuracy feedback after each training trial, and the passing criterion was set to 80% accuracy in their decisions on which of the two intervals (the test or reference interval) was longer. The number of training trials was initially set to 10, but increased automatically by another 10 trials if participants failed to pass the criterion.

## Results and Discussion

**Fig. A2.**
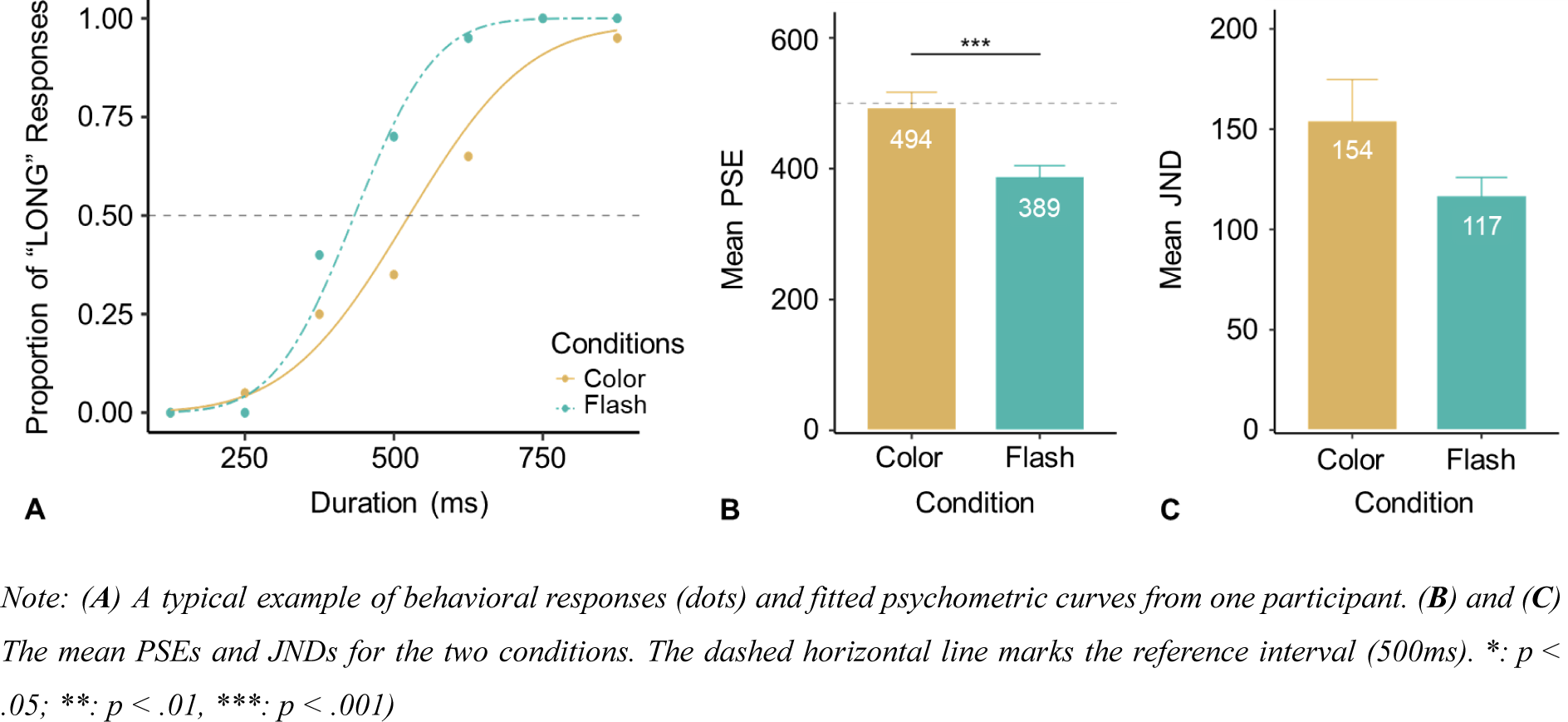
Results from Experiment 2.

In order to ensure high-quality data, participants’ performance was considered valid only if they were able to identify 125 ms as ‘short’ and 875 ms as ‘long’ (in comparison with the 500-ms reference interval) above 75% correct. Out of the 20 participants, only 3 did not meet this criterion and so their data were excluded from further analysis. Fig A2 shows the proportion of ‘long’ responses and the associated psychometric curves from a typical participant.

The mean PSE (with associated standard error) was 494 (± 23) for the Color condition and 389 (± 16) ms for the Flash condition. A *t*-test revealed the PSE to be significantly lower (105 ms) in the flash vs. the color condition, *t*(16) = 4.95, *p* < .001. This suggests that time distortions caused by the attentional blink were more pronounced due to the flashes as compared to the color changes. This also confirms the finding of Experiment 1, where the PSE (in the No-Gap/Fixation condition) was shorter than the reference duration of 500-ms. The mean JNDs (± SE) were 154 (± 20) and 117 (± 9) for the Color and Flash conditions, respectively; the difference was non-significant, *t*(16) = 1.74, *p* = .102, indicative of comparable discrimination difficulty.

To summarize, the results of the control experiment showed that replacing flashes with feature (color) changes to establish intervals significantly reduced time distortion caused by the attentional blink. In fact, with color changes defining the intervals, the onset bias was effectively eliminated, as the PSE (494 ms) did not differ from the 500-ms standard (*t*(16) = −0.28, *p* = .786). Given this, we used color changes to define the target and reference intervals in Experiment 2.

## Footnotes

1 Participants who failed to pass the accuracy criterion after two rounds of training, as well as those whose eyes could not be reliably tracked by the Eyelink system (e.g., because they wore glasses), did not proceed to the formal experiment. Also, a number of participants exerted their right to quit the experiment without completing the total number of trials, stating mainly the demandingness of the task and tiredness as a reason. They were all nevertheless paid for their service at the standard rate. Overall, this led to the loss of 7 participants in Experiment 1 and 8 in Experiment 2.

2 Given this, the non-test interval could potentially also have been used as the reference interval. This is, however, unlikely, as participants were instructed (and in the practice trials trained) to compare the two like-colored – test and reference – intervals.

3 In any case, our observed underestimation of the first interval would also be in line with (Morrone et al., 2005; Yarrow et al., 2001)) finding of saccadic compression rather than Chronostasis.

